# Bactericidal efficiency and mechanism of specifically targeted antimicrobial peptides optimized based on structural and functional relationships

**DOI:** 10.1101/679977

**Authors:** Peng Tan, Zhenheng Lai, Yongjie Zhu, Changxuan Shao, Muhammad Usman Akhtar, Weifen Li, Xin Zheng, Anshan Shan

**Author notes:** Corresponding author: Anshan Shan, Postal address: No. 600 Changjiang Road, Xiangfang District, Harbin, China. These authors contributed equally to this work (Peng Tan, Zhenheng Lai).

## Abstract

In contrast to traditional broad-spectrum antibiotics, it is difficult for bacteria to develop resistance to most specifically targeted antimicrobial peptides (STAMPs), moreover, they can maintain a normal ecological balance and provide long-term protection for the body. However, therapeutic applications of STAMPS are hindered by their weak activity, and imperfect specificity as well as lack of knowledge to understand their structure-activity relationships. To further investigate the effects of different parameters on the biological activities of STAMPs, a peptide sequence, WKKIWK^D^PGIKKWIK, was truncated, extended, and provided with an increased charge and altered amphipathicity. In addition, a novel template modification method was introduced, in which a phage-displayed peptide that recognized and bound to *E. coli* cells was attached at the end of the sequence. Compared with the traditional template modification method, peptide 11, which contained a phage-displayed peptide at the C-terminus, exhibited superior narrow-spectrum antibacterial activity against *E. coli* compared to that of parental peptide 2, and the activity and specificity of 11 were increased by 5.0 and 2.4 times, respectively. Additionally, 11 showed low cell toxicity and relatively desirable salt, serum, acid and alkaline stability. In this study, 11 specifically killed *E. coli* by causing cytoplasmic membrane rupture and cytosol leakage. In summary, these findings are useful for improving the activity and specificity of STAMPs and show that peptide 11 is better able to combat the growing threat of *E. coli* infections.

## Introduction

With the development of bacterial resistance, antibiotics to combat microbes are becoming limited. Moreover, antibiotics are used to kill bacteria and result in the release of large amounts of lipopolysaccharide (LPS), which is the major component of the outer membrane of Gram-negative bacteria and induces serious complications, such as septic shock and severe sepsis[1]. For that reason, there is a urge need for development of novel antimicrobial agents. Antimicrobial peptides (AMPs) are a new class of antimicrobial agent that has the potential to substitute for traditional antibiotics.

Antimicrobial peptides (AMPs), which are part of the immune system, form the first line of defense against pathogenic bacteria[2]. Compared with traditional antibiotics, AMPs possess a unique mechanism, and it is widely accepted that the cytoplasmic membrane is the main target of AMPs[3]. Because of this mechanism, bacteria have low potential to develop resistance to AMPs. However, AMPs have similar activity profile to conventional antibiotics, showing broad-spectrum antimicrobial activities against both Gram-negative and Gram-positive bacteria; these agents kill or inhibit benign and pathogenic organisms indiscriminately, thus disrupting the homeostasis between a healthy microbiota and the immune system[4, 5]. Consequently, there is an urgent need for novel antimicrobial peptides that are capable of targeting specific pathogens without harming the normal flora. Specifically targeted antimicrobial peptides (STAMPs) can maintain the ecological balance of normal microbial communities[6]. Although researchers are now working on the development of targeted antibacterial agents, some problems persist, such as the weak activity and poor specificity of STAMPs. At present, the strategies used to improve the activity of broad-spectrum antimicrobial peptides are systematic sequence extension or truncation, amino acid substitution, and increases in charge and amphipathicity[7], but how to improve the activity and specificity of STAMPs has been rarely reported to date.

Recently, we discovered the antimicrobial peptide KI[8]. After removing the chemical modification at both ends, we found that it only had weak antibacterial activity against *Escherichia coli*(*E. coli*). As is known, *E. coli* is the predominant facultative flora of the human and animal intestine. Pathogenic *E. coli* not only can cause bacterial diarrhea in animals but can also lead to human urinary tract infections, meningitis, and pneumonia and even affect the human nervous system[9, 10]. To further improve the activity and specificity of peptide 2 for *E. coli* and to study the relationship between the structure and activity of STAMPs, we modified peptide 2 using traditional template modification methods, including altering its length, amphipathicity and charge[11, 12]. In addition, we used a novel modification method to link a phage-displayed peptide to the ends of the peptide 2. Filamentous bacteriophages can display foreign peptides expressed by DNA sequences introduced in the genome through recombinant modification on their surface, which are able to recognize and bind specific targets, such as the whole *E. coli* cell or some kinds of cell surface receptors[13, 14]. In this study, we used phage displayed-peptide 14 screened from a phage display random peptide library, which targets *E. coli* cell and specifically binds to the surface of these cells.

The secondary structures of these peptides were determined in different solutions (PBS, SDS, and TFE). The antimicrobial activities of the peptides were then measured, and we introduced a novel index, the targeted antimicrobial index (TI), which can reflect the specificity of STAMP for a kind of bacteria. Moreover, the salt, serum acid and alkaline sensitivities; hemolytic activity; and cytotoxicity were also evaluated. Finally, to study the antibacterial mechanism of STAMP, LPS and LTA binding, outer membrane permeability, inner membrane permeability, cytoplasmic membrane depolarization, scanning electron microscopy (SEM), transmission electron microscopy (TEM), super-resolution microscopy (SRM), flow cytometry were also employed. We found that linking a phage-displayed peptide to the C- and N-terminus could significantly improve the antimicrobial activity and specificity of STAMP. Concurrently, we designed peptides 1-10 using different traditional sequence modification methods, but their effects were worse than our novel proposed method. These findings are helpful for the development of design strategies based on STAMPs.

## Results

### Peptide characterization

The designed peptides were synthesized by Sangon Biotech (Shanghai, China) and purified to greater than 95% purity using analytical reverse phase high-performance liquid chromatography (RP-HPLC). The molecular weights of peptides were determined by matrix-assisted laser desorption/ionization time-of-flight mass spectroscopy (MALDI-TOF MS). The measured molecular weights were close to the theoretical molecular weights, which indicated that the peptides were successfully synthesized.

The following design methods were adopted to modify the parental peptide 2. 1) Change the length of parental peptide 2. The ends of parental peptide 2 were truncated and extended, and each peptide was simultaneously decreased or increased by two amino acids at both ends. Among them, peptide 1 was decreased by a total of 4 amino acids and peptides 3 and 4 were centered on the ^D^PG Angle of peptide 2 and sequentially extended at the N- and C-terminus in the order of KWIK and IKKW, respectively. Peptides 3 and 4 were increased by a total of 4 and 8 amino acids, respectively. 2) Increase charge of the parental peptide 2. To peptides 5 and 6 were added one and two lysines at the end of the parental peptide, respectively. 3) Enhance the amphipathicity of parental peptide 2. As shown in Figure 1A, peptides 7 and 8 were designed by substituting amino acids at the 2nd or 13th position of parental peptide 2. Peptide 9 was designed by swapping the position of the 2nd and 13th amino acids of parent peptide 2. In peptide 10, the β-turn unit was removed to become a perfect amphipathic peptide. 4) Parental peptide 2, which connects the phage-displayed peptide at the N- or C-terminus, was used to enhance the specific identification and binding ability of STAMP to *E. coli*. Phage-displayed peptide 14, which was recruited from random peptide libraries, a whole-cell phage-display method was used to isolate peptide 14 specifically binding to the cell surface of *E. coli*[15]. Moreover, during the construction of the novel antimicrobial peptides with multi-domains, space hindrance often occurred because the different domains of the peptides were too close to each other, thus inhibiting the biological functions of the different peptide segments[6]. In the design of the peptide sequences, when the amino acid sequence was added to one end of the parental peptide, we introduced a short peptide sequence, GGG, composed of a flexible amino acid G as a linker[6]. The amino acid sequences of the peptides are listed in Table 1. The wheel-diagram, 3D structure projection, schematic structure and schematic model as shown in Figure 1.

**Figure 1.**
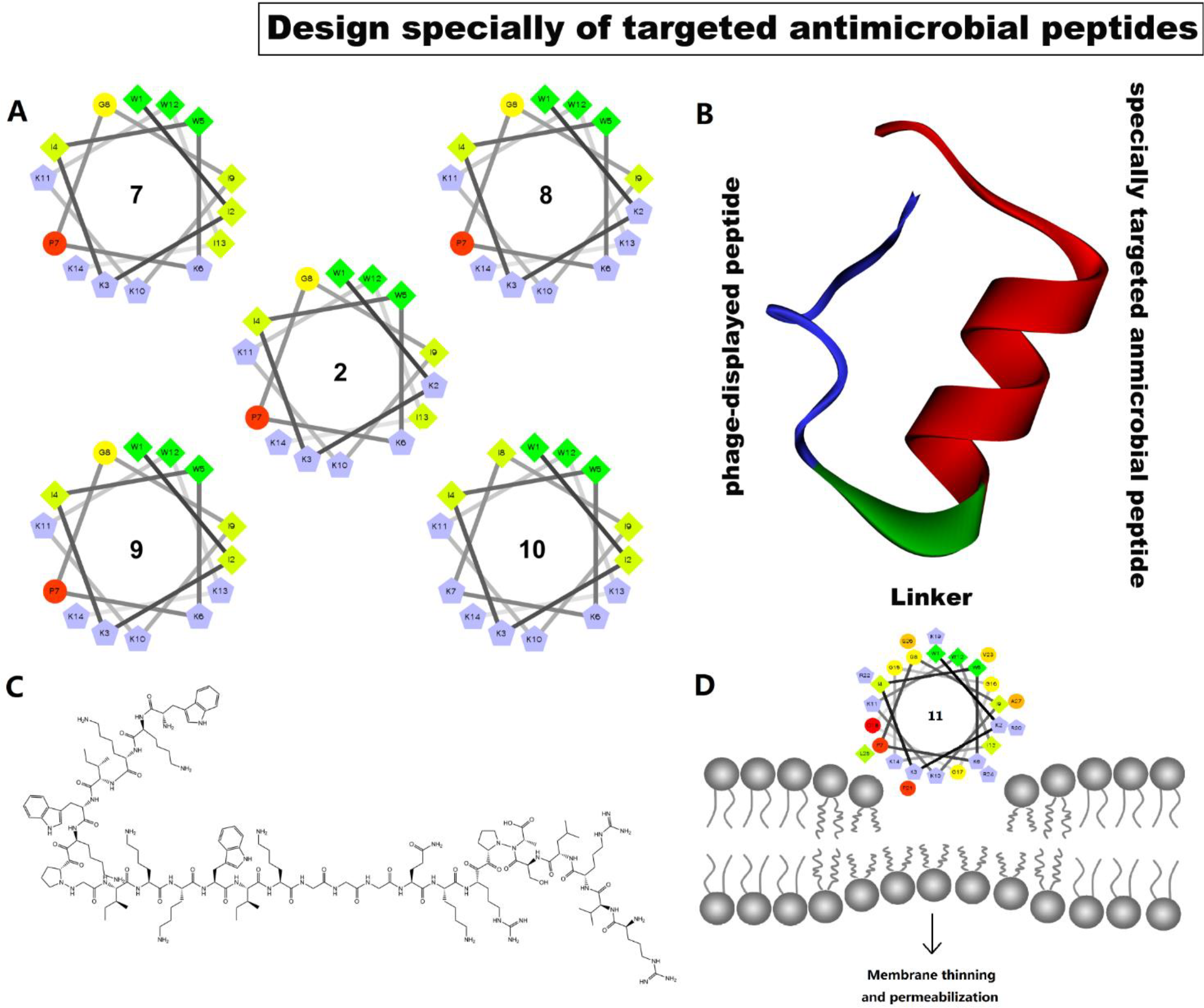
Design of synthetic specially targeted antimicrobial peptides. (A) Helical wheel projections of the peptides. By default the output presents the hydrophilic residues as circles, hydrophobic residues as diamonds, potentially negatively charged as triangles, and potentially positively charged as pentagons. Hydrophobicity is color coded as well: the most hydrophobic residue is green, and the amount of green is decreasing proportionally to the hydrophobicity, with zero hydrophobicity coded as yellow. Hydrophilic residues are coded red with pure red being the most hydrophilic (uncharged) residue, and the amount of red decreasing proportionally to the hydrophilicity. The potentially charged residues are light blue. (B) Three-dimensional structure projections of the 11. The STAMP domain, linker and displaying peptide are color coded as red, green and blue. (C) Sequence and schematic structure of the 11. (D) Schematic model of the interaction of 11 with *E. coli* membrane.

**Table 1.**
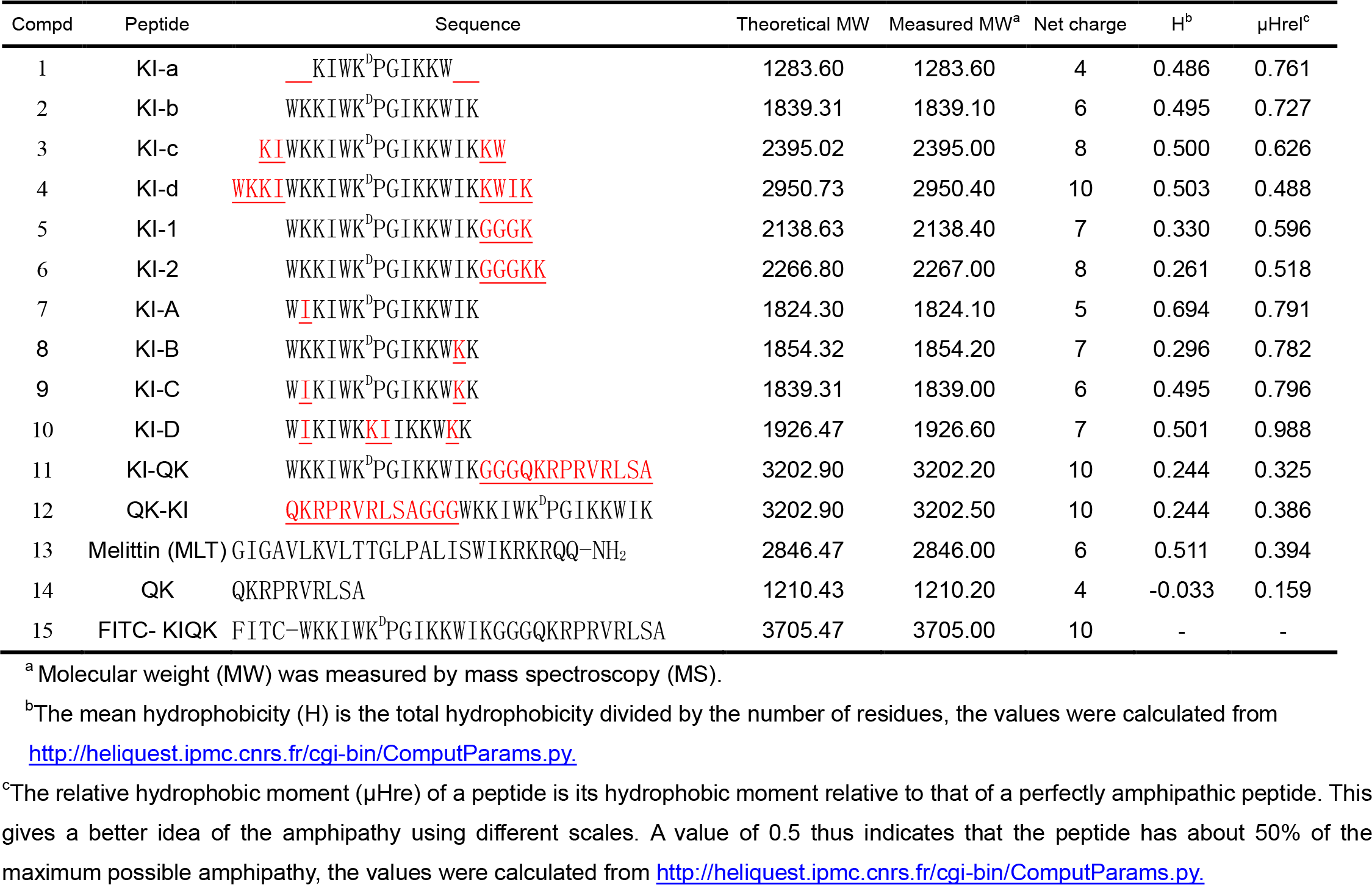
Peptide Design and Their Key Physicochemical Parameters. Changed amino acids are shown in red font and underlined.

### CD spectra

The secondary structures of the peptides were investigated by CD spectroscopy. SDS micelles were used to simulate the anionic microbial membrane environment, and TFE was used to mimic the hydrophobic environment[16]. As shown in Figure 2, the spectra of most peptides were not characteristic of the helix conformation in 10 mM PBS. By contrast, in 50% TFE, all peptides other than 14 tended to form an α-helical structure with two negative peaks at approximately 208 and 222 nm. In the SDS environment, the [θ]220 values of peptide 1 (−728), 2 (−839), 3 (−1668) and 4 (−6924) showed that when ^D^PG was in the middle of the peptide sequence, the weaker helical structure of the peptide was destroyed by ^D^PG with the extension of the peptide sequence. Peptide 5 and 6 also showed helical properties in SDS environments. The amphiphilicity of peptide 7, 8 and 9 was improved; however, the helical propensity changes were not obvious. When ^D^PG was removed, peptide 10 was converted into a typical α-helical structure. Phage-displayed peptide 14 had a minimum at ≈ 200 nm and a value near zero at 220 nm, supporting a disordered structure, and 11 and 12 exhibited two negative dichroic bands at approximately 208 and 222 nm, indicating the predominance of α-helical structures, similar to the results obtained for parental peptide 2.

**Figure 2.**
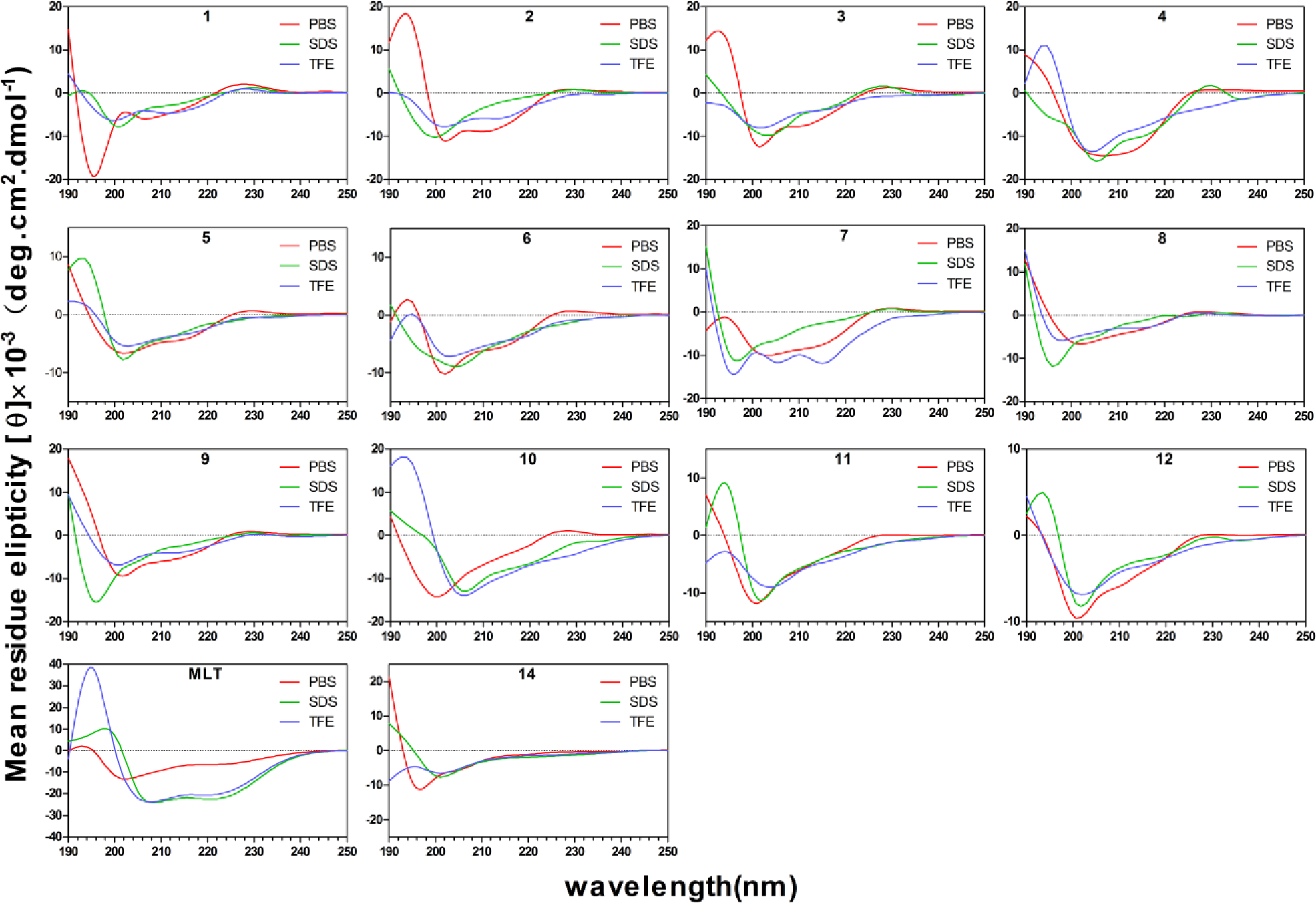
CD spectra of all peptides. All the peptides were dissolved in 10 mM PBS (PH 7.4), 50% TFE or 30 mM SDS. The mean residual ellipticity was plotted against wavelength. The values from three scans were average per sample, and peptide concentration were fixed at 150 μM. The spectra were smoothed by GraphPad software.

### Antibacterial activity and specificity of peptides

Antibacterial activity of peptides is summarized in Table 2. To better evaluate the antimicrobial activity and specificity of peptides against *E. coli*, the geometric means (GMs) of the MICs of the peptides against the tested pathogenic or beneficial strains were calculated and are presented in Table 3. In the design of truncated and extended STAMP chains, stronger anti-*E. coli* activity was achieved with a longer length of the peptide (peptide 1 < 2 < 3 < 4). The increased charge of peptides 5 and 6 did not show a significant enhancement of antimicrobial activity against *E. coli* compared with parental peptide 2 (peptide 2 ≈ 5 ≈ 6). In the design of the amphipathicity modification, the activity of the peptide against *E. coli* showed a decreasing trend. However, when STAMP was replaced with perfect amphipathic peptide 10, the activity of peptide 10 toward *E. coli* increased (peptide 7 ≈ 8 ≈ 9 < 2 < 10). When the phage-displayed peptide was linked to the end of the parental peptide, the activities of peptides 11 and 12 to *E. coli* were enhanced; 11 had better antimicrobial activity against *E. coli* (peptide 2 < 12 < 11). Among all peptides, 4 and 10 displayed broad-spectrum anti-bacterial activity; in addition, the other peptides showed narrow-spectrum antimicrobial activity against *E. coli*. In the MIC test for beneficial bacteria, melittin possessed broad-spectrum antimicrobial activity, and no other peptides except peptide 4 had any effect. Moreover, the ratio of GM (*E. coli*) to GM (other pathogenic strains) and GM (*E. coli*) to GM (beneficial strains) was used to as the targeting index (TI); smaller targeting index values indicated greater specific antimicrobial ability toward *E. coli*. Peptide 11 had the lowest TI_all_ value (0.028), which was 31 times lower than that of melittin (TI_all_=0.876). Compared with parental peptide 2 (TI_all_=0.067), the specificity of 11 for *E. coli* increased by 2.4 times; the specificities of 3 (TI_all_=0.040) and 12 (TI_all_=0.035) were also enhanced.

**Table 2.** MICs^a^(μM) of the Engineered Peptides against pathogenic or beneficial strains.

| | Minimum inhibitory concentration (MIC) in μM | | | | | | | | | | | | | |
| --- | --- | --- | --- | --- | --- | --- | --- | --- | --- | --- | --- | --- | --- | --- |
|  | 1 | 2 | 3 | 4 | 5 | 6 | 7 | 8 | 9 | 10 | 11 | 12 | MLT | 14 |
| Pathogenic strains |  |  |  |  |  |  |  |  |  |  |  |  |  |  |
| *E. coli* 25922 | >128 | 16 | 4 | 2 | 8 | 8 | 32 | 16 | 16 | 8 | 2 | 8 | 4 | 64 |
| *E. coli* UB1005 | >128 | 16 | 2 | 1 | 4 | 4 | 16 | 8 | 8 | 8 | 2 | 8 | 2 | 128 |
| *E. coli* K88 | >128 | 8 | 2 | 0.5 | 8 | 8 | 16 | 16 | 16 | 8 | 4 | 4 | 2 | 64 |
| *E. coli* K99 | >128 | 16 | 2 | 1 | 16 | 32 | 16 | 32 | 16 | 4 | 4 | 8 | 2 | 64 |
| *E. coli* 078 | >128 | 32 | 8 | 2 | 64 | 64 | 128 | 64 | 128 | 16 | 8 | 32 | 4 | >128 |
| *E. coli* 987P | >128 | 16 | 2 | 1 | 8 | 8 | 16 | 16 | 8 | 2 | 2 | 4 | 2 | 128 |
| *S. aureus* 29213 | >128 | >128 | 32 | 4 | >128 | >128 | >128 | >128 | >128 | 4 | >128 | >128 | 4 | >128 |
| *S. aureus* 25923 | >128 | >128 | 32 | 4 | >128 | >128 | >128 | >128 | >128 | 8 | >128 | >128 | 4 | >128 |
| *S. epidermidis* 12228 | >128 | >128 | 16 | 2 | >128 | >128 | 128 | >128 | >128 | 4 | >128 | 128 | 4 | >128 |
| *P. aeruginosa* 27853 | >128 | >128 | 16 | 2 | >128 | >128 | >128 | >128 | >128 | 4 | 64 | >128 | 4 | >128 |
| *S. typhimurium* 7731 | >128 | 128 | 32 | 2 | 128 | 128 | >128 | >128 | 64 | 128 | 64 | 128 | 4 | >128 |
| *S. agalactiae* 13813 | >128 | >128 | >128 | 2 | >128 | >128 | >128 | >128 | >128 | 8 | >128 | >128 | 4 | >128 |
| Beneficial strains |  |  |  |  |  |  |  |  |  |  |  |  |  |  |
| *L. plantarum* 8014 | >128 | >128 | >128 | 64 | >128 | >128 | >128 | >128 | >128 | 128 | >128 | >128 | 2 | >128 |
| *L. rhamnosus* 7469 | >128 | >128 | >128 | 64 | >128 | >128 | >128 | >128 | >128 | 128 | >128 | >128 | 2 | >128 |
| *L. rhamnosus* 1.9205 | >128 | >128 | >128 | 32 | >128 | >128 | >128 | >128 | 128 | >128 | >128 | >128 | 2 | >128 |
| *L. rhamnosus* 1.9011 | >128 | >128 | >128 | 64 | >128 | >128 | >128 | >128 | 128 | >128 | >128 | >128 | 2 | >128 |
| *L. rhamnosus* 1.0385 | >128 | >128 | >128 | 32 | >128 | >128 | >128 | >128 | 64 | 128 | >128 | >128 | 2 | >128 |
| *L. rhamnosus* 1.0386 | >128 | >128 | >128 | 64 | >128 | >128 | >128 | >128 | >128 | >128 | >128 | >128 | 4 | >128 |

<sup>a</sup> Minimum inhibitory concentration (MIC, μM) was determined as the lowest concentration of peptide that inhibited 95% of the bacterial growth. Data are representative of three independent experiments.

**Table 3.** MHC, GM, SI and TI Values of the Engineered Peptides.

| Peptide | MHC<sup>a</sup> | GM<sup>b</sup> |  |  | SI<sup>d</sup> | TI<sup>e</sup> |  |  |
| --- | --- | --- | --- | --- | --- | --- | --- | --- |
|  |  | *E. coli* | Pathogenic strains<sup>c</sup> | Beneficial strains | *E. coli* | Pathogenic strains<sup>c</sup> | Beneficial strains | All |
| 1 | >128 | 256.000 | 256.000 | 256.000 | 1.000 | 1.000 | 1.000 | 1.000 |
| 2 | >128 | 16.000 | 228.070 | 256.000 | 16.000 | 0.070 | 0.063 | 0.067 |
| 3 | >128 | 2.828 | 35.919 | 256.000 | 90.523 | 0.079 | 0.011 | 0.040 |
| 4 | 16 | 1.122 | 2.520 | 50.797 | 14.260 | 0.445 | 0.022 | 0.233 |
| 5 | >128 | 11.314 | 228.070 | 256.000 | 22.627 | 0.050 | 0.044 | 0.047 |
| 6 | >128 | 12.699 | 228.070 | 256.000 | 20.159 | 0.056 | 0.050 | 0.053 |
| 7 | >128 | 25.398 | 228.070 | 256.000 | 10.080 | 0.111 | 0.099 | 0.105 |
| 8 | >128 | 20.159 | 256.000 | 256.000 | 12.699 | 0.079 | 0.079 | 0.079 |
| 9 | >128 | 17.959 | 203.187 | 161.270 | 14.255 | 0.088 | 0.111 | 0.100 |
| 10 | >128 | 6.350 | 8.980 | 181.019 | 40.315 | 0.707 | 0.040 | 0.376 |
| 11 | >128 | 3.175 | 161.270 | 256.000 | 80.630 | 0.020 | 0.035 | 0.028 |
| 12 | >128 | 8.000 | 203.187 | 256.000 | 32.000 | 0.039 | 0.031 | 0.035 |
| MLT | 2 | 2.520 | 4.000 | 2.245 | 0.794 | 0.630 | 1.122 | 0.876 |
| 14 | >128 | 101.594 | 256.000 | 256.000 | 2.520 | 0.397 | 0.397 | 0.397 |

<sup>a</sup>MHC is the minimum hemolytic concentration that caused 5% hemolysis of human red blood cells. When no detectable hemolytic activity was observed at 128 μM, a value of 256 μM was used to calculate the selectivity index.
<sup>b</sup>The geometric mean (GM) of the peptide MICs against bacteria was calculated. When no detectable antimicrobial activity was observed at 128 μM, a value of 256 μM was used to calculate the geometric mean.
<sup>c</sup>Pathogenic strains here refer to bacteria other than *E. coli*.
<sup>d</sup>SI is calculated as MHC/GM (*E. coli*). Larger values indicate greater cell selectivity.
<sup>e</sup>The targeting index represents the specific ability to specially target antibacterial peptides. TI is calculated as GM(*E. coli*)/GM(Other Pathogenic strains) and GM(*E. coli*)/GM(Beneficial strains). Smaller values indicate greater specific ability.

### Hemolytic activity and cytotoxicity

Table 3 and Figure 3 summarize the peptide hemolytic activities. The results showed that hemolysis of all peptides except 4 was less than 5% at all concentrations. Compared to melittin, all peptides had significantly lower hemolytic activity (P<0.05). The ratio of MHC to GM served as the selectivity index (SI), which indicated the cell selectivity of a peptide. Peptide 11 was found to have a relatively high SI of 80.630, and although peptide 3 had a higher selectivity index than peptide 11, its TI was only 0.040. In the cytotoxicity test of human embryonic kidney cells (HEK293T) and intestinal porcine enterocyte cells (IPEC-J2) (Figure 4), the toxicity of extended peptide 4 and perfect amphipathic peptide 10 was significant; peptide 4 killed approximately 99% of HEK293t and IPEC-J2 at 128 μM, followed by peptide 10, which killed approximately 99% of HEK293t and 63% IPEC-J2. In contrast, 11 exhibited very high selectivity for two kinds of cells, and compared with 2, the toxicity was increased minimally at 128 μM.

**Figure 3.**
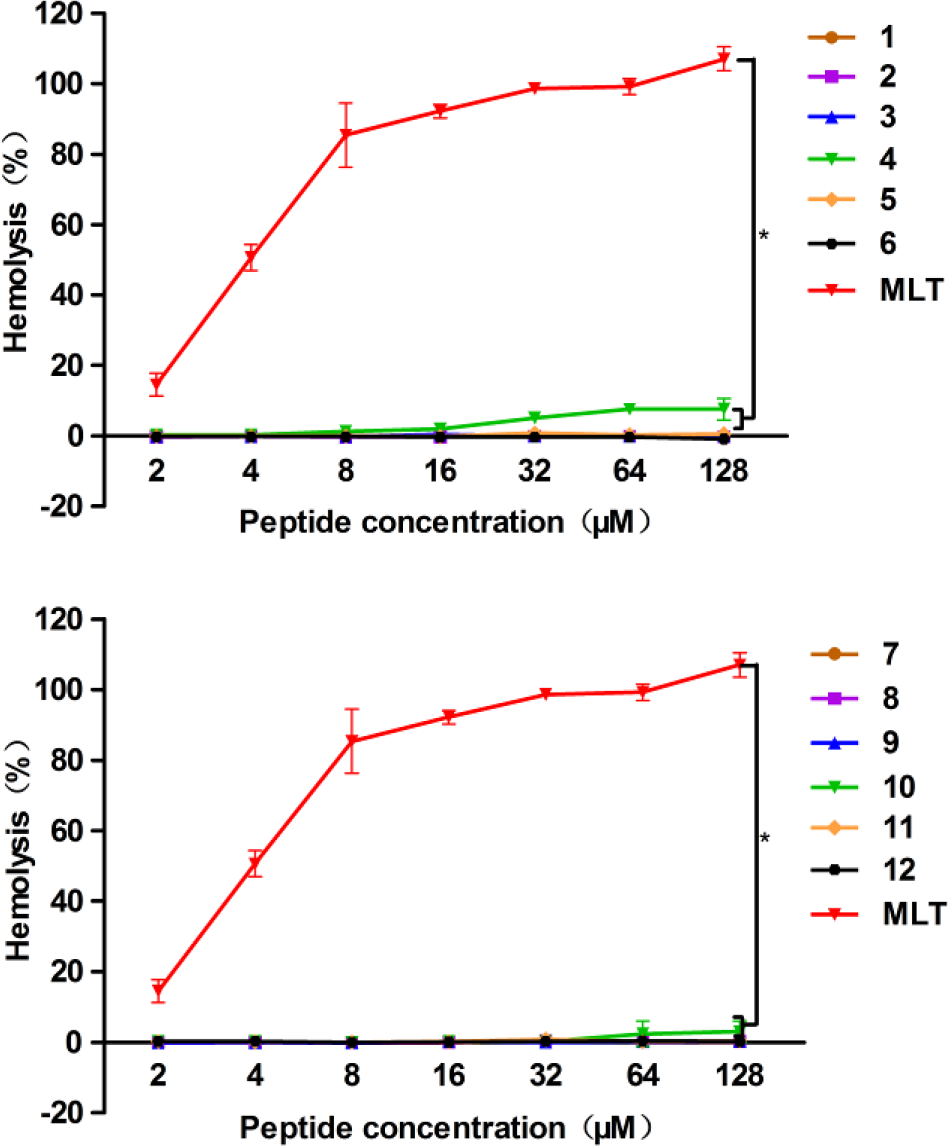
Hemolytic activity of the peptides against hRBCs. The graphs were derived from the average of three independent trials:(*) P < 0.05, compared to values for Melittin.

**Figure 4.**
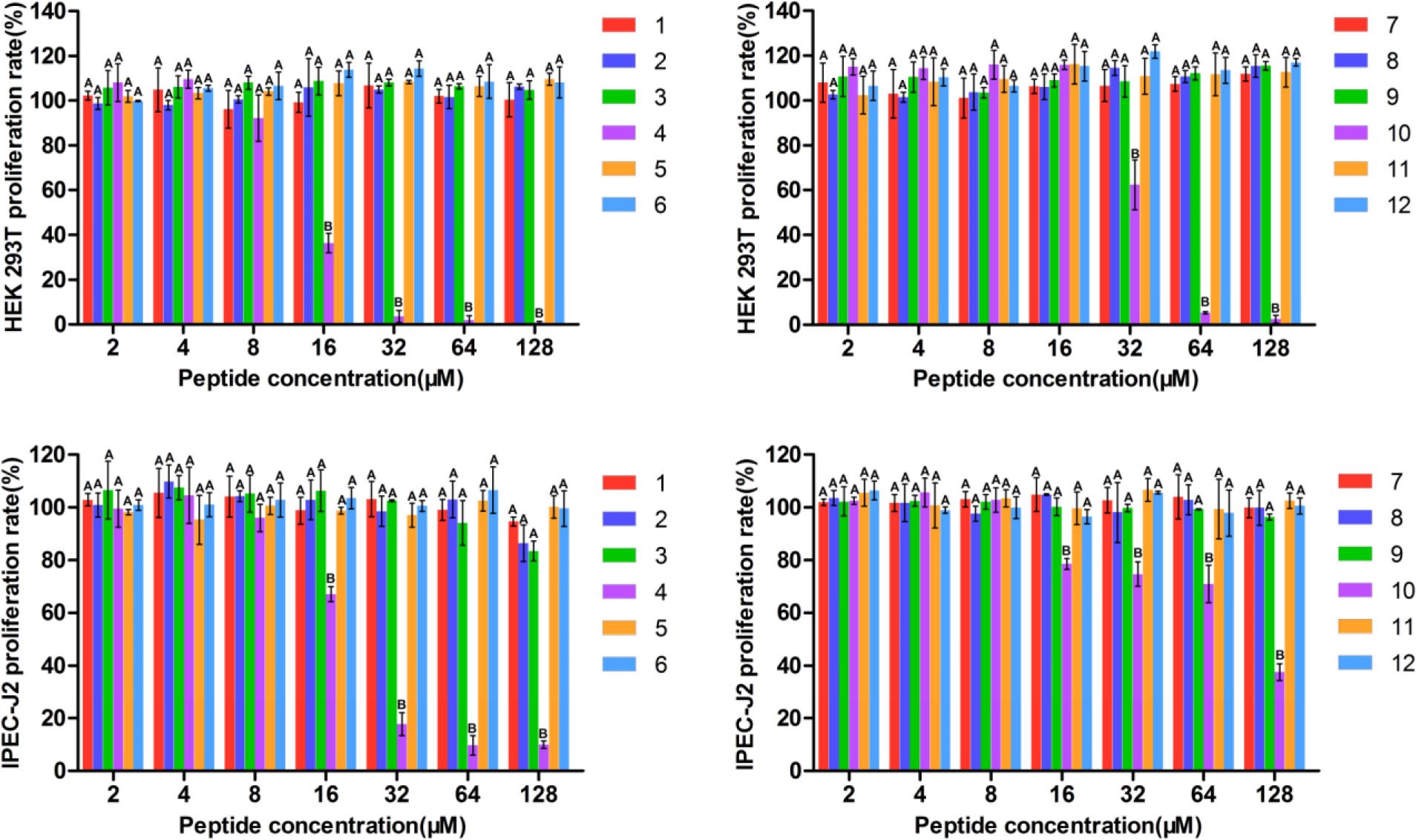
Cytotoxicity of the peptides against HEK293T, IPEC-J2 cells. The graphs were derived from average of three independent trials. Mean values in the same concentration with different superscript indicate a significant difference (P < 0.05).

### Salt, Serum, Acid and Alkali sensitivity

Peptides must maintain activity in a physiological environment for clinical applications. However, many peptides have low stability *in vivo* and are susceptible to salt, serum, acid and alkaline conditions. Thus, the antimicrobial activities of the peptides were tested following the addition of concentrations of different salts, serum, acid and alkaline agents using a sensitivity assay (Table 4). The results revealed that in the presence of 150 mM NaCl and 1 mM MgCl_2_, these peptides (other than 4) significantly reduced the potency against *E. coli* 25922. Parental 2 almost lost all antibacterial activity toward *E. coli*. Nonetheless, peptide 11, which we selected as a target sequence, was not completely deprived of activity, maintaining a relatively desirable active state. The MIC values for *E. coli* were 32 μM and 16 μM. Overall, 2 showed greater susceptibility compared with 11.

**Table 4.** MIC Values of the Engineered Peptides against *E. coli* ATCC25922 in the Presence of Physiological Salts, Serum, Acid and Alkaline.

| Peptide | Control<sup>a</sup> | Physiological Salts |  |  |  |  |  | Serum |  |  | Acid and Alkaline |  |  |  |
| --- | --- | --- | --- | --- | --- | --- | --- | --- | --- | --- | --- | --- | --- | --- |
|  |  | NaCl<sup>b</sup> | KCl<sup>b</sup> | MgCl<sub>2</sub><sup>b</sup> | ZnCl<sub>2</sub><sup>b</sup> | NH<sub>4</sub>Cl<sup>b</sup> | FeCl<sub>3</sub><sup>b</sup> | 50% | 25% | 12.5% | PH=2 | PH=4 | PH=10 | PH=12 |
| 1 | >64 | >64 | >64 | >64 | >64 | >64 | >64 | >64 | >64 | >64 | >64 | >64 | >64 | >64 |
| 2 | 16 | >64 | 16 | 32 | 16 | 16 | 16 | 16 | 16 | 16 | 32 | 16 | 16 | 32 |
| 3 | 4 | 16 | 4 | 8 | 4 | 4 | 4 | 4 | 4 | 4 | 4 | 4 | 4 | 4 |
| 4 | 2 | 2 | 2 | 2 | 2 | 2 | 2 | 2 | 4 | 2 | 2 | 2 | 2 | 2 |
| 5 | 8 | >64 | 8 | 32 | 8 | 8 | 8 | 8 | 8 | 8 | 32 | 32 | 16 | 16 |
| 6 | 8 | >64 | 8 | 32 | 8 | 8 | 8 | 8 | 8 | 8 | 32 | 16 | 16 | 16 |
| 7 | 32 | >64 | 32 | >64 | 32 | 32 | 32 | 32 | 32 | 32 | 32 | 32 | 32 | 32 |
| 8 | 16 | >64 | 16 | 32 | 16 | 16 | 16 | 16 | 16 | 16 | 64 | 32 | 16 | 64 |
| 9 | 16 | >64 | 32 | 64 | 16 | 16 | 16 | 32 | 16 | 16 | 32 | 32 | 16 | 16 |
| 10 | 8 | 32 | 8 | 64 | 8 | 8 | 8 | 8 | 8 | 8 | 8 | 8 | 8 | 8 |
| 11 | 2 | 32 | 2 | 16 | 2 | 4 | 4 | 2 | 2 | 2 | 4 | 4 | 2 | 4 |
| 12 | 8 | >64 | 8 | 32 | 8 | 8 | 8 | 4 | 4 | 4 | 16 | 8 | 16 | 16 |
| MLT | 4 | 16 | 4 | 8 | 4 | 4 | 4 | 4 | 4 | 4 | 4 | 4 | 4 | 4 |
| 14 | 64 | >64 | >64 | >64 | >64 | >64 | >64 | >64 | >64 | 64 | 64 | >64 | >64 | 64 |

<sup>a</sup>The control MIC values were determined in the absence of these physiological salts, serum, acid and alkaline.
<sup>b</sup>The final concentrations of NaCl, KCl, NH<sub>4</sub>Cl, MgCl<sub>2</sub>, ZnCl<sub>2</sub>, and FeCl<sub>3</sub> were 150 mM, 4.5 mM, 1 mM, 8μM, 6μM and 4μM.

In the presence of NH_4_Cl, KCl, ZnCl_2_ and FeCl_3_ displayed relatively minimally repressive effects on the anti-bacterial activities of the peptides. In serum, acid and alkaline sensitivity tests, the MIC values of the peptides were not significantly increased.

### LPS, LTA binding assays

Lipopolysaccharide (LPS) and lipoteichoic acid (LTA) are negative electrochemical components of Gram-negative and Gram-positive bacteria, respectively, which can interact with positively charged AMPs. As shown in Figure 5A and Figure 5B, the binding activity of 11 to LPS and LTA presented a dose-dependent increase, the binding capacity of which was stronger than that of 2 and comparable to that of melittin. Figure 5A and Figure 5B show a stronger fluorescence intensity for the binding activity of 11 with 50% *E. coli* at a low peptide concentration (2 μM) than that of melittin (21%) and parental 2 (6%), and the binding capacity of 11 to *E. coli* LPS at 2 μM was stronger than that of *P. aeruginosa* LPS (16%) and *S. aureus* LTA (23%).

**Figure 5.**
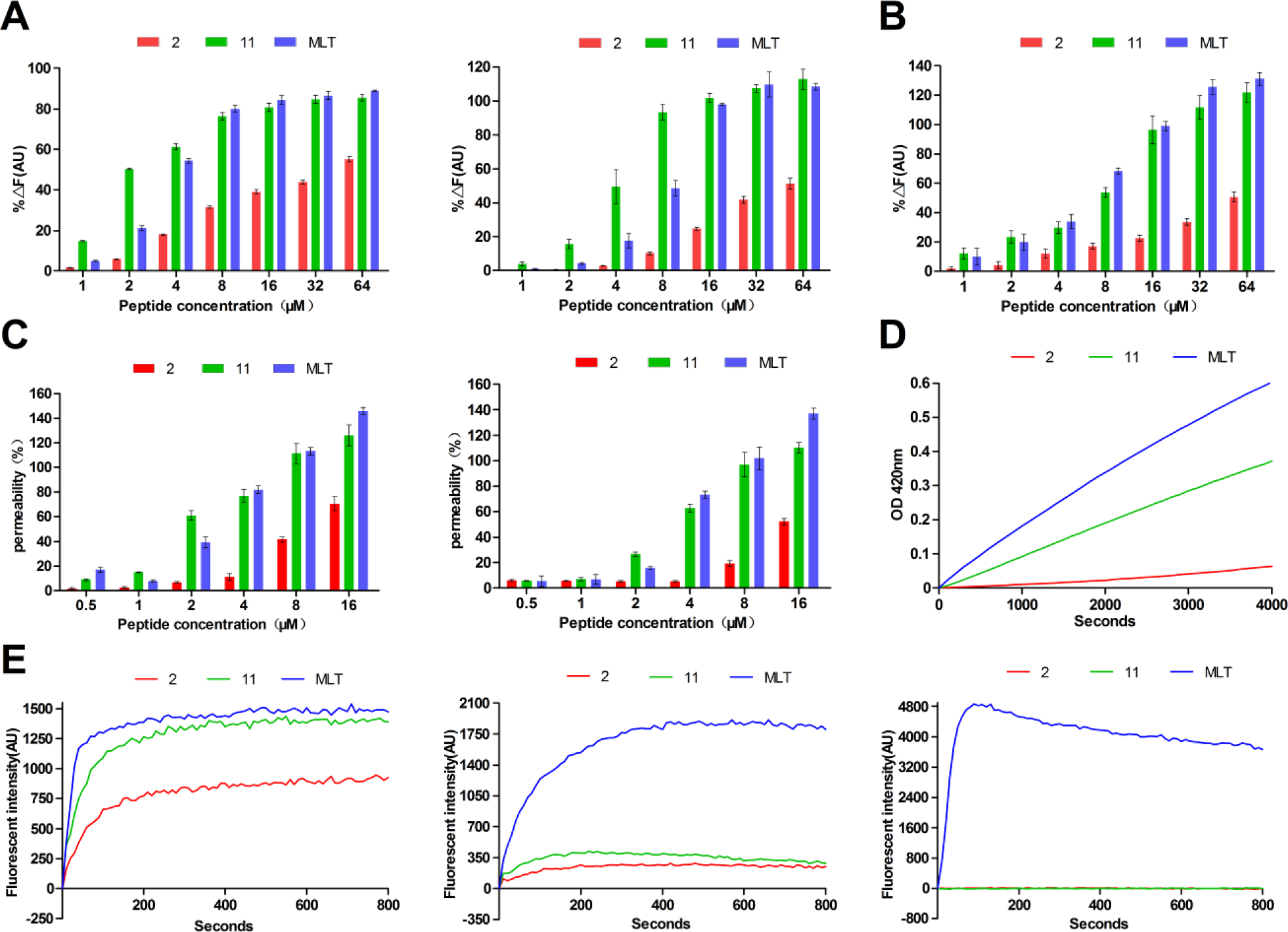
(A) Peptide binding affinity to LPS from *E. coli* 0111:B4 and *Pseudomonas aeruginosa* 10. (B) Peptide binding affinity to LTA from *S. aureus.* (C) Outer membrane permeability of the *E. coli* 25922 and *P. aeruginosa* 27853 treated by 2 μM peptides. (D) Inner membrane permeability of the *E. coli* 25922 treated by 2 μM peptides. (E) The cytoplasmic membrane potential variation of *E. coli* 25922, *P. aeruginosa* 27853 and *S. aureus* 29213 treated by 2 μM peptides.

### Outer and inner membrane permeability

The potential of the peptides to permeabilize bacterial outer membranes was studied using *N*-phenyl-1-naphthylamine (NPN) uptake assays[17]. The outer membrane is a unique component of Gram-negative bacteria, which can provide an extra layer of protection to the organism. When the outer membrane is damaged or disrupted, NPN binds to the phospholipid layer and provide fluorescence. As shown in Figure 5C, a dose-dependent response of peptides in permeabilizing the outer membrane of Gram-negative bacteria (*E. coli* and *P. aeruginosa*) was observed. In the presence of 2 μM peptide 11, the permeability of the outer membrane was stronger than that of parental 2 and melittin, reaching 61% and 27%, respectively, for *E. coli* and *P. aeruginosa.*

The inner membrane permeability was assessed by o-nitrophenol-β-D-galactoside (ONPG) assays, which were used to detect the enterobacter in delayed lactose fermentation (containing Escherichia, Klebsiella, etc.)[18]. When the inner membrane of *E. coli* is damaged, ONPG enters the cytoplasm, where it can be hydrolyzed into galactose and yellow o-nitrophenol (ONP) by beta-galactosidase. Therefore, inner membrane permeabilization can be measured by color changes in the culture medium. As shown in Figure 5D, peptides 2 and 11 and melittin induced increases in the permeability of the inner membrane from 0 to 4000 seconds at 2 μM. The optical density of 11 was significantly higher than that of 2 and similar to that of melittin, indicating that the inner membrane permeability of 11 was obviously stronger than that of parental 2. This phenomenon was largely consistent with the results of the outer membrane permeability test.

### Cytoplasmic membrane depolarization

The ability of 2, 11 and melittin to depolarize the membranes of intact *E. coli, P. aeruginosa* and *S. aureus* cells was determined using the membrane potential-sensitive dye diSC_3_-5, which can become concentrated in the cytoplasmic membrane based on the membrane potential, leading to the self-quenching of fluorescence[19]. When the membrane potential is disrupted, the dye dissociates into the buffer, causing an increase in fluorescence intensity. As shown in Figure 5E, in the cytoplasmic membrane test of *E. coli*, the depolarization ability of 11 was enhanced compared with that of parental 2. By contrast, 2 and 11 showed slight depolarization of the cytoplasmic membrane of *P. aeruginosa* compared with melittin, potentially causing a small number of strains to rupture. For *S. aureus*, no cytoplasmic membrane depolarization occurred within 800 seconds, suggesting that 2 and 11 were not effective against membranes of *S. aureus*. We consider the above results to explain the narrow-spectrum antibacterial activity of the peptides against *E. coli.*

### SEM and TEM characterization

To further elucidate the narrow-spectrum antimicrobial mechanisms of 11, SEM and TEM were implemented to study *E. coli, P. aeruginosa* and *S. aureus* morphological alterations, and parental 2 and melittin were added for comparison. To evaluate the therapeutic effect of the peptides, we used the same concentration of 2 μM. In the absence of peptide treatment, the bacterial cells had brilliant and smooth membrane surfaces (Figure 6). However, one hour after treatment with 11, membrane creping and destruction were observed in *E. coli* cells; however, this phenomenon was not observed in most *P. aeruginosa* and *S. aureus* and cells. In contrast to 11, parental 2 did not achieve an antibacterial effect due to an insufficient concentration, and the *E. coli* cells were barely damaged. Melittin showed a broad-spectrum antibacterial effect at this concentration, with significant membrane stunting and corrugation on the surface of *E. coli, P. aeruginosa* and *S. aureus*.

**Figure 6.**
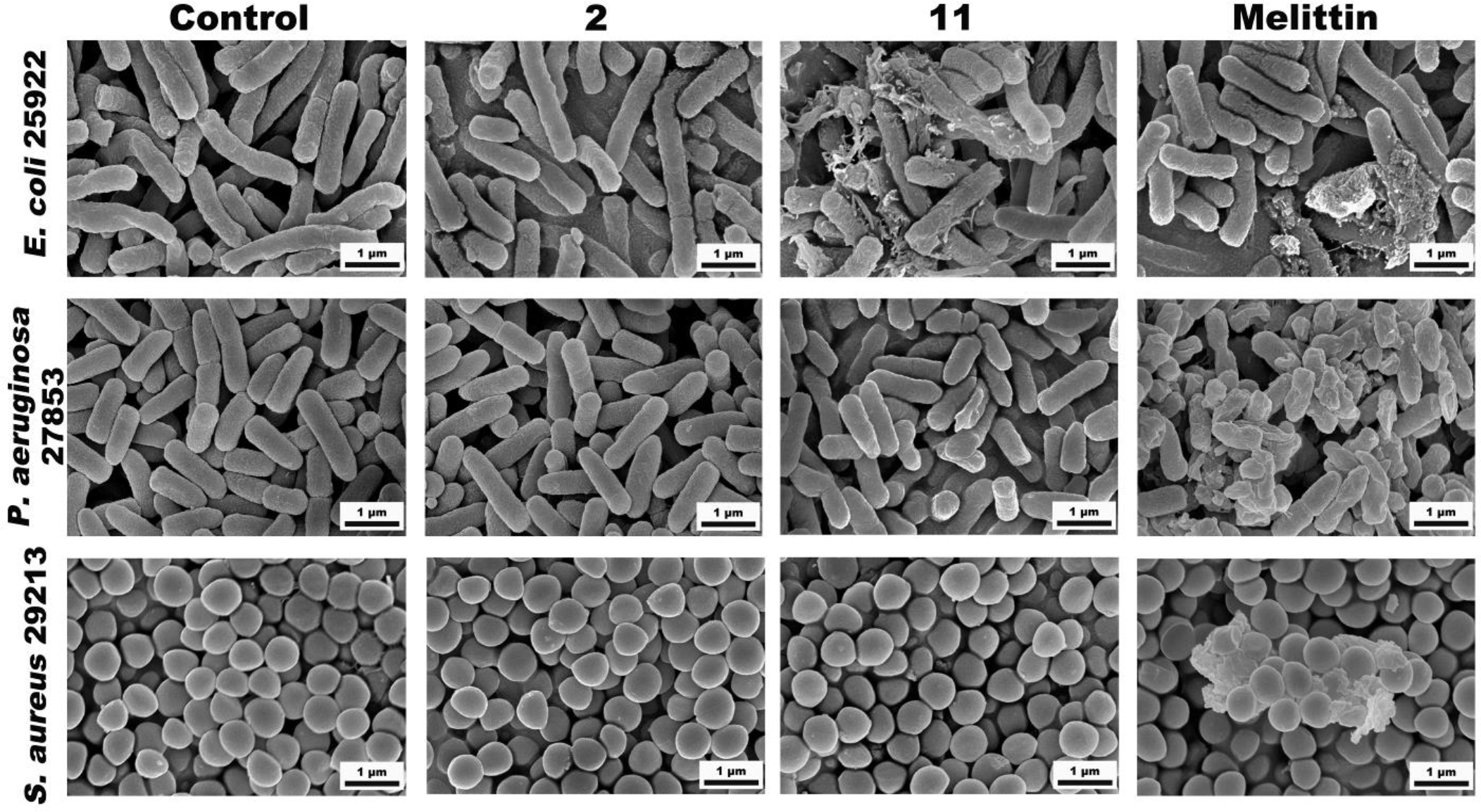
SEM images of *E. coli* 25922, *P. aeruginosa* 27853 and *S. aureus* 29213 treated for 1 h with the 2 μM peptides and 10 mM PBS (pH 7.4) (control).

TEM revealed the morphological and intracellular changes in *E. coli, P. aeruginosa* and *S. aureus* cells after treatment with 2, 11 and melittin. Similarly, under TEM, after 60 min of treatment, 11 caused substantial damage to *E. coli* membranes and outflow of the cytoplasm (Figure 7).

**Figure 7.**
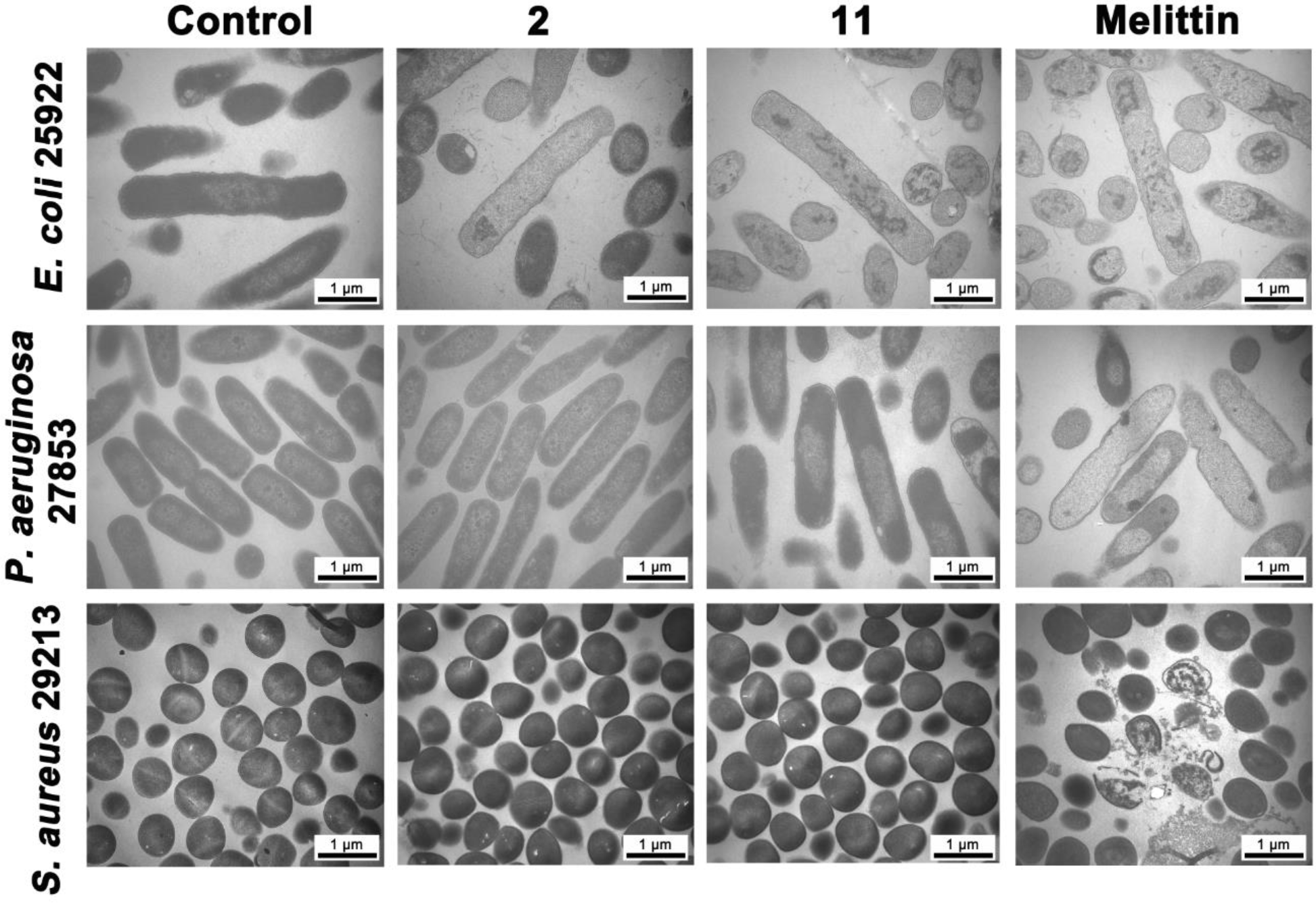
TEM images of *E. coli* 25922, *P. aeruginosa* 27853 and *S. aureus* 29213 treated for 1 h with the 2 μM peptides and 10 mM PBS (pH 7.4) (control).

### Super-resolution microscopy (SRM)

The localization of FITC-labeled peptide 11 was studied by super-resolution microscopy. As shown in Figure 8, FITC-labeled peptide 11 was represented by green fluorescent signals on the surface and inside of *E. coli* cells, while *E. coli* nucleic acid was stained with PI dye and observed as a red fluorescent signal. The results distinctly indicated that 11 targeted the *E. coli* membrane surface and compromised membrane integrity.

**Figure 8.**
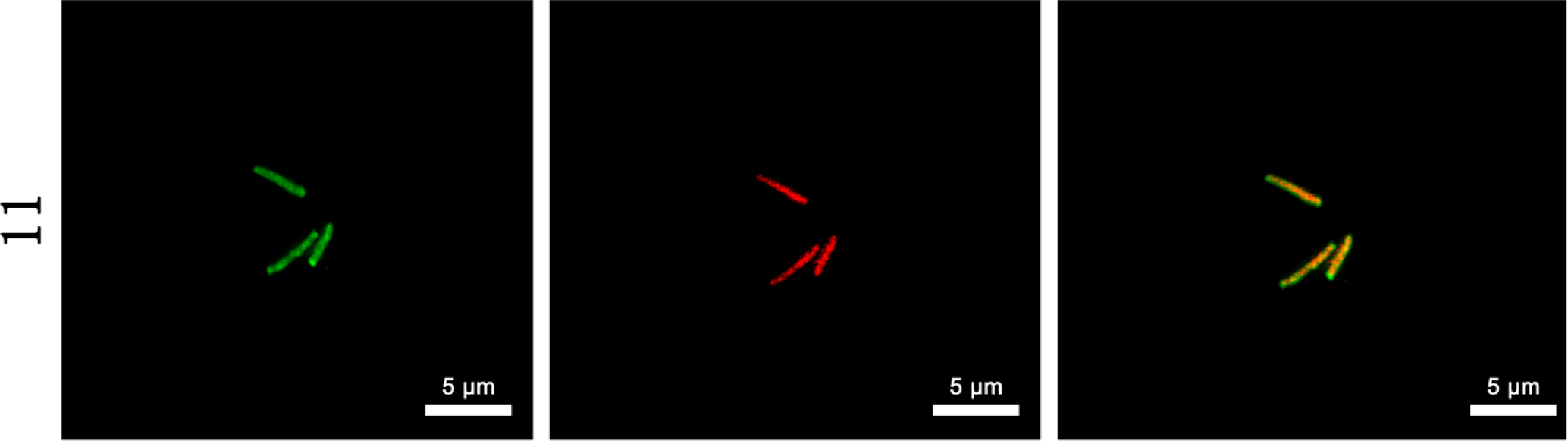
Deltavision OMX system analysis of *E. coli* 25922. Green signal from FITC-11 peptide, red signal from PI.

### Flow cytometry

The DNA intercalating dye propidium iodide (PI) was used as an indicator to investigate *E. coli, P. aeruginosa* and *S. aureus* cell death by flow cytometry (Figure 9). In the absence of peptide, the percentage of *E. coli* 25922, *P. aeruginosa* 27853 and *S. aureus* 29213 cells with PI fluorescent signal was 2.3%, 5.0%, and 5.2%, respectively. Treatment with 2 μM peptides showed that peptide 2 resulted in 57.7% (*E. coli*), 31.3% (*P. aeruginosa*), and 14.9% (*S. aureus*) cell staining; peptide 11 resulted in 96.2% (*E. coli*), 33.0% (*P. aeruginosa*), and 11.0% (*S. aureus*) cell staining; and melittin resulted in the staining of greater than 90% staining of all three bacteria. These results indicated that peptide 11 had narrow-spectrum antibacterial activity against *E. coli* and induced more potent damage to *E. coli* than the parental peptide.

**Figure 9.**
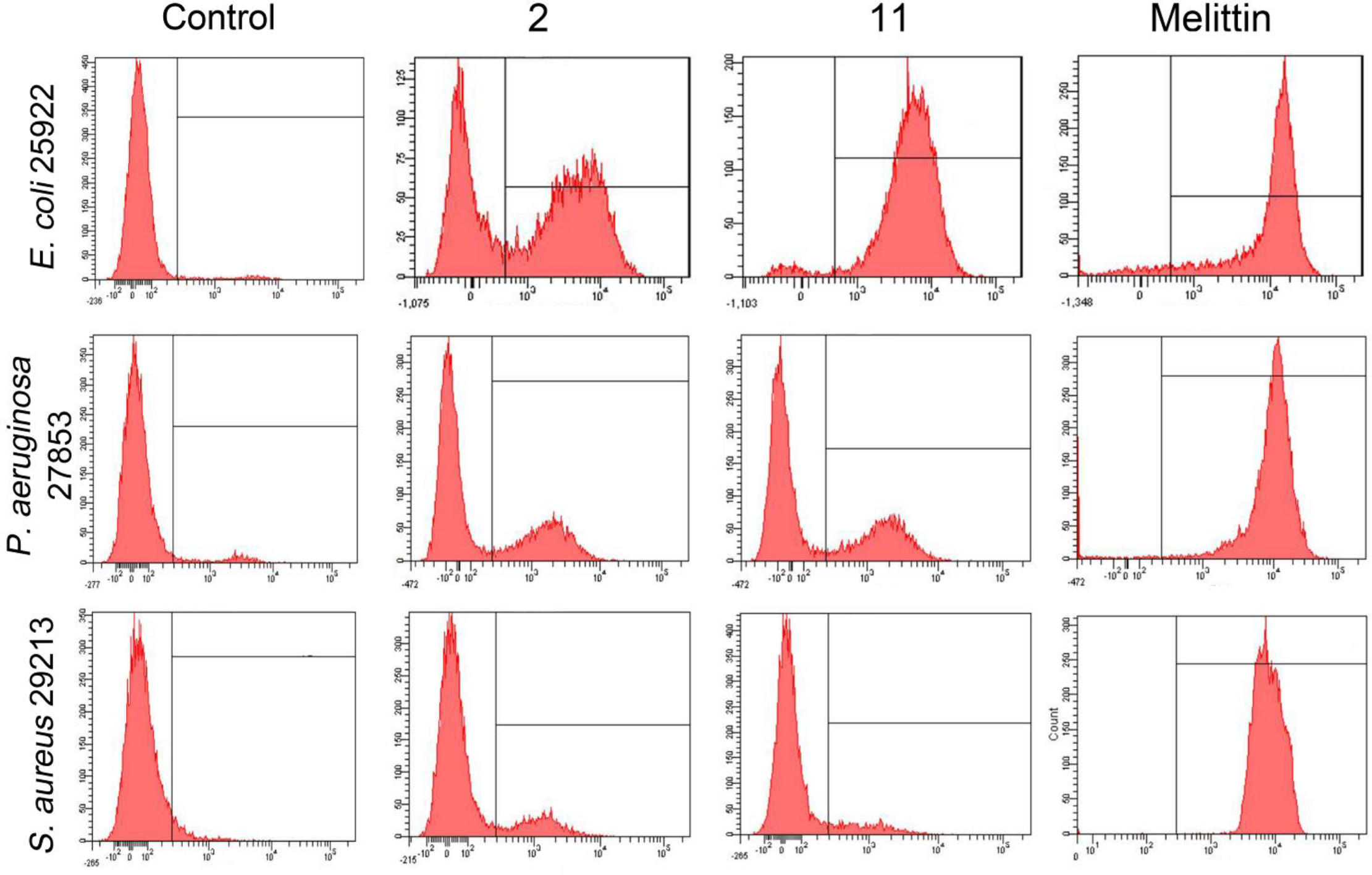
Flow cytometric analysis of *E. coli* 25922, *P. aeruginosa* 27853, and *S. aureus* 29213. The increments of cellular fluorescence intensity of PI (10μg/mL) after treating with the peptides was analyzed by flow cytometry.

## Discussion

With the increase in antibiotic-resistant bacteria worldwide, the demand for novel antimicrobial drugs to fight against infectious diseases is also increasing[20]. STAMPs represent a novel class of therapeutic drugs that can be used to treat microbial infections without disrupting homeostasis. However, there are many obstacles in the development of STAMPs that must be circumvented, including optimization of STAMP specificity and activity. Template modification is performed using a naturally occurring or designed AMP sequence as a starting template, followed by systematic sequence extension or truncation, a charge increase, amino acid substitutions, and amphipathicity changes to improve antimicrobial activities[7]. Previous researchers have commonly changed one or two key parameters to transform peptides, resulting in few optimal sequences for therapeutic applications due to weak activity; susceptibility to salt; serum, acid, and alkaline inactivation; and high cytotoxicity. In this study, all the above factors were used in an attempt to improve the activity and specificity of STAMP, and we focused on the effects of different design parameters on the antimicrobial spectrum and biological activity of peptides.

The stable secondary structure of AMPs in the membrane environment is very important for their biological activity[21]. Our results showed that most of the AMPs with disordered conformations in PBS displayed a conversion to an α-helical conformation in TFE and SDS (Figure 2). The helical propensity of AMPs is typically affected by the composition of hydrophobic amino acids. Phage-displayed peptide 14 contained only three hydrophobic amino acids as well as proline, and proline residues can damage the α-helical propensity. Thus, peptide 14 tended to form a disordered structure in membrane-like environments. Similarly, D-Pro-Gly had a rigid bend that also disrupted the α-helical propensity; therefore, except peptide 10, these peptides had a weak α-helical secondary structure signal in a membrane-like environment. In comparison, peptide 10 and melittin had the typical α-helical conformation.

As is known, the length of the sequence affects the activity of AMPs[22, 23]. We designed peptide 1 by removing two amino acids at both the N- and C-termini of parental peptide 2 without changing the amino acid composition, which resulted in a loss of biological activity. This phenomenon suggested that shortening the length of peptide 2 could not achieve the desired effect and that reducing four or more reside(s) led to a gradual loss of biological activity. Moreover, peptide 1 had a reduced helical tendency in SDS (Figure 2), suggesting that STAMPs require at least 14 amino acids to maintain helical a structure able to span the membrane lipid bilayer. To further determine the optimal length of the peptide for exerting narrow-spectrum antibacterial activity against *E. coli*, peptide 2 was extended, and the designed peptides 3 and 4 showed increased activity against *E. coli*. Interestingly, peptide 4, which contained 22 amino acids, exhibited broad-spectrum antibacterial activity, and the results showed that STAMP could be converted into a broad-spectrum antimicrobial peptide after extending the sequences. Although the activity of peptide 3 and 4 was enhanced in comparison to those of parental peptide 2, the increase in sequence length also increased the toxicity of AMPs[24]. This phenomenon is consistent with previous studies and might provide an explanation for the excess hydrophobicity of the longer peptide molecule to obstruct the interaction between the peptide and zwitterionic phospholipids while leading to a loss of cell selectivity[25, 26]. Therefore, in this study, we did not enhance the specificity and activity of STAMPs by extending the sequence because this design approach conflicts with the low cytotoxicity (Figures 3 and 4).

The bacterial membrane contains anionic components, such as lipopolysaccharides, mannoproteins, or anionic phospholipids, which can interact with AMP molecules via electrostatic interactions[27]. AMPs aggregate on the membrane surface when they accumulate at a critical concentration, followed by membrane lipid bilayer insertion, resulting in the destruction of the membrane, leakage of the cytoplasm and cell death[28, 29]. Therefore, we speculate that increasing the charge number of STAMPs can enhance the electrostatic interactions between the molecule and the membrane to improve the antimicrobial activity and specificity. Arginine and lysine act as cationic residues that can produce strong antibacterial activity under physiological conditions, while the use of arginine residues is frequently associated with relatively higher hemolytic activities[30, 31]. To avoid increased toxicity, we added one or two lysine residues to the C-terminus of parental peptide 2 and designed peptides 5 and 6. Unfortunately, the activities of peptides 5 and 6 toward *E. coli* were not enhanced compared with that of peptide 2 (Table 2 and Table 3). Therefore, these findings indicate that STAMPs are the same as broad-spectrum antimicrobial peptides, and the relationship between the charge and biological activity is not linear, as increases of positive charges above a threshold (usually 5-6) did not result in increased antimicrobial activity[26, 32].

Amphipathicity is another important parameter that can affect the antimicrobial activity of α-helical AMPs[33]. The research of Khara, J. S showed that a perfect amphipathic of an α-helical AMP can effectively enhance the antibacterial potential, but perfect amphipathicity often leads to a simultaneous increase in activity and cytotoxicity[34]. Thus, a balance of amphipathicity is required to achieve the satisfactory biological activity. In our search, peptides 7, 8, 9 and 10 contained amino acid replacements to increase amphipathicity, but the results showed that the antibacterial activity of peptides 7, 8, and 9 against *E. coli* was weaker than that of parental peptide 2. In 30 mM SDS, the helical structures of peptides 7, 8, and 9 were not significantly affected by the enhanced amphipathicity (Figure 2); therefore, the activity of the peptides was not improved. Simultaneously, perfect amphipathic peptide 10 showed broad-spectrum antibacterial activity, as well as increased hemolysis and cytotoxicity. This result further confirmed that perfect amphipathicity often results in increased hemolysis and cytotoxicity[35, 36]. Moreover, this phenomenon showed that the antimicrobial activity and specificity of STAMPs could not be enhanced by simply increasing the amphipathicity and that increasing amphipathicity may lead to a transformation of STAMPs into broad-spectrum antimicrobial peptides.

Although the above structural parameters did not supply the STAMPs with ideal antibacterial activity, a delicate balance among them may be required for the design of an ideal STAMP with increased antibacterial activity and specificity. Therefore, hybrid peptides have become an attractive method to optimize these sequences[37]. In this study, we used a phage-displayed peptide that was capable of specific and strong binding to *E. coli* cells[15]. This peptide was attached to the N- or C-termini of parental peptide 2 to give peptide 11 and 12 respectively. With the fusion of 14, 11 and 12 displayed 5.0 and 2.0 times increased antimicrobial activity relative to that of parental peptide 2 alone. A theory to explain this phenomenon might be that 11 and 12 have a better hydrophobic and hydrophilic phase balance than the parental peptide, resulting in enhanced antimicrobial activity because positive residues and hydrophobic residues can bind to and insert into the cytoplasmic membranes[35]. Moreover, the specific recognition ability of 11 and 12 to *E. coli* was enhanced by the addition of the phage-displayed peptide. Hemolysis and cytotoxicity are important factors that limit the clinical application of antimicrobial peptides; thus, it is important to evaluate the cytotoxic effect of peptides. Fortunately, our target peptide 11 showed lower hemolysis and cytotoxicity (Figure 3 and Figure 4) at the level of the average *E. coli* MIC value, demonstrating that the selectivity of 11 for the *E. coli* cell membrane exceeded that for mammalian cell membrane[23].

Previous reports have shown that the activity and cytotoxicity of peptides are often correlated with their hydrophobicity and helical tendency[38, 39]. In membrane-like environments, 11 exhibited α-helical characteristics and had weaker hydrophobicity and helical tendencies than melittin, potentially indicating why target peptide 11 showed antimicrobial activity against *E. coli* and low cytotoxicity. This result further confirmed that the activity and cytotoxicity were associated with the hydrophobicity and helical tendencies. Therefore, when designing hybrid peptides using a phage-displayed peptide, the optimum balance of hydrophobicity and helical tendency must be maintained.

Positively charged salt ions compete with peptide molecules to reduce the antibacterial activity of AMPs[38]. In our research, Na^+^ and Mg^2+^ compromised the antimicrobial activity of peptides other than 4 against *E. coli* (Table 4). At physiological concentrations, Na^+^ and Mg^2+^ reduce interactions between AMPs and the membrane via the charge-shielding effect and thus reduce the antimicrobial activity against *E. coli*. Moreover, Mg^2+^ can bind LPS on the outer membrane of *E. coli* to prevent the proximity of antibacterial peptide molecules[41, 42]. Therefore, increasing the charge can effectively replace the salt ions around the membrane by an ion-exchange mechanism. Our results showed that the increased charge of peptide 11 via terminal link phage-displayed peptide led to a reduction of salt ion sensitivity. Furthermore, previous studies have confirmed that increases in hydrophobicity can also reduce the adverse effects of salt ions on the antimicrobial activity of AMPs[43]. Peptide 4 maintained broad-spectrum antimicrobial activity at a physiological salt concentration, which might be related to both the overall hydrophobicity and the increased charge. Compared with our target peptide 11, peptide 4 showed the same amount of charge as peptide 11 (+10), but the overall hydrophobicity was stronger than 11; therefore, peptide 4 showed improved salt stability. Moreover, all the designed peptides could retain their original biological activities in serum, acid and alkaline environments. In conclusion, our target peptide 11 maintained relatively satisfactory activity in a complex physiological environment compared with its parental peptide 2.

Previous studies have proposed that broad-spectrum AMPs exert antimicrobial activity by membrane permeabilization[44, 45]. However, the bactericidal mechanism of STAMPs is still unclear. Therefore, we added the Gram-negative bacteria *P. aeruginosa* and Gram-positive bacteria *S. aureus* as controls because previous MIC tests showed that peptide 11 did not have the ability to kill these two strains (Table 2). LPS and LTA are the main components of Gram-negative bacteria and Gram-positive bacteria, respectively, which can interact with AMPs via electrostatic interactions[44]. In our study, the binding affinity of peptide 11 to LPS and LTA showed a dose-dependent effect (Figure 5A and Figure 5B), which demonstrated that the interactions between STAMP 11 and LPS and LTA were only electrostatic and that there was no specificity between strains, so peptide 11 could accumulate on the surface of the Gram-negative and Gram-positive bacterial membrane. The outer membrane is a unique component of Gram-negative bacteria, and AMP molecules must penetrate the outer membrane to get close to the cytoplasmic membrane[47]. In general, the outer membrane of all Gram-negative bacteria possesses a certain degree of permeability, allowing the exchange of small molecules and ions between the cell interior and the extracellular medium[48, 49]. Peptide 11 could enhance the outer membrane permeability of *E. coli* and *P. aeruginosa* in a dose-dependent manner (Figure 5C), but the penetration of the outer membrane was not sufficient to kill the strains. When AMPs penetrate the outer membrane and local peptides reach the threshold, AMP molecules can insert their hydrophobic cores into the phospholipid bilayer of the cytoplasmic membrane, disrupting the membrane surface potential and forming a mass of pore channels, finally resulting in cell lysis[50]. In cytoplasmic membrane depolarization assays, we demonstrated that 11 could perturb the cytoplasmic membrane potential of *E. coli*, had a slight effect on *P. aeruginosa*, but had no effect on *S. aureus* (Figure 5E). Therefore, we believe selective destruction of the cytoplasmic membrane of *E. coli* by 11 supports a narrow-spectrum antibacterial activity of the peptide against these cells (Figure 10). The inner membrane permeability results further indicated that 11 could induce cytoplasmic membrane leakage (Figure 5D). Direct observation by SEM and TEM further confirmed the membrane damage effects of 11 on *E. coli.* The super-resolution microscopy results showed that 11 damaged the cell membrane of *E. coli.* Furthermore, flow cytometry analysis indicated that linkage of the displayed peptide at the end of the parental peptide improved its anti-bacterial activity against *E. coli.* Altogether, 11 was observed to penetrate the outer membrane of Gram-negative bacteria and destroy the cytoplasmic membrane potential to induce the release of the cell contents, eventually leading to the death of *E. coli.*

**Figure 10.**
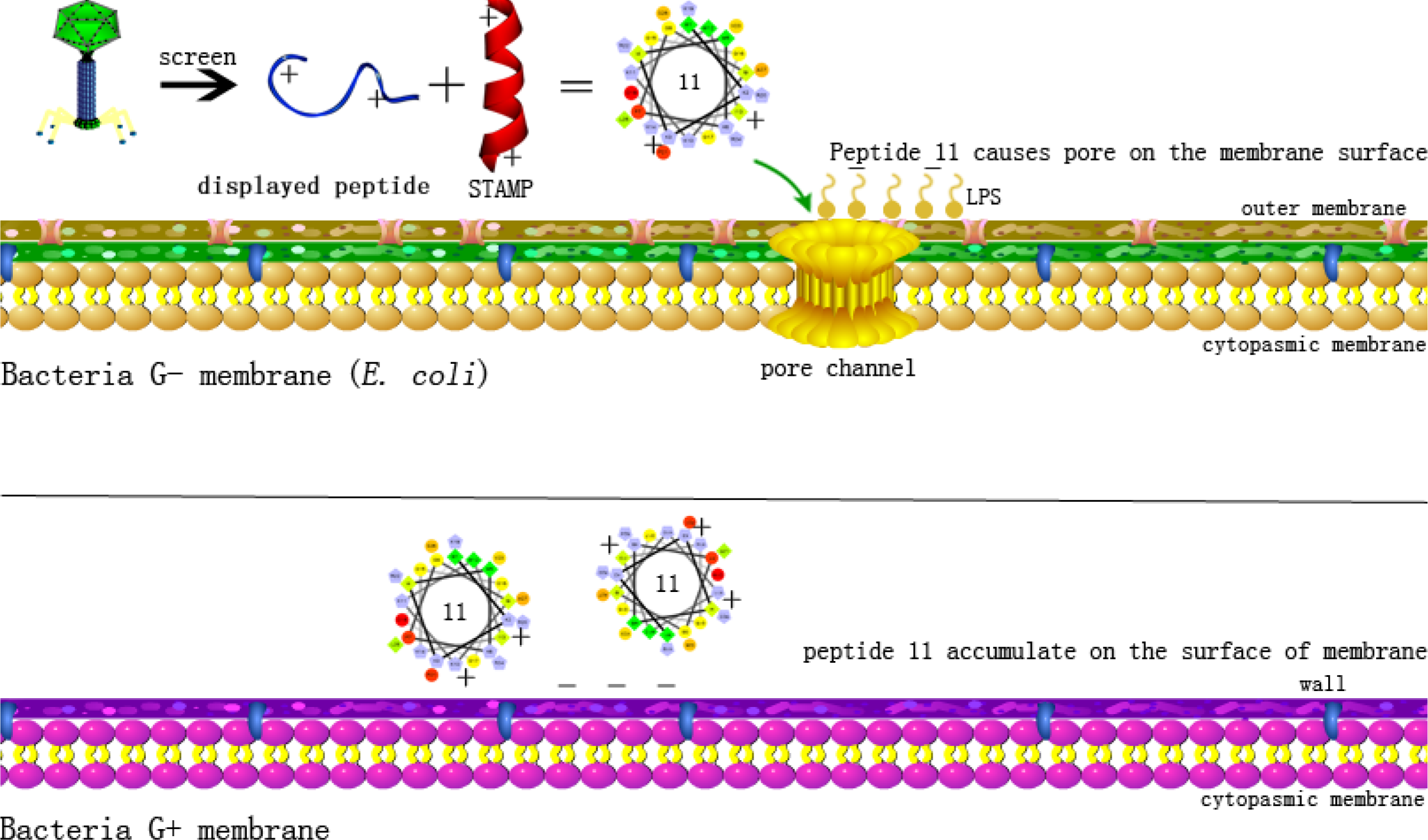
Schematic diagram of bactericidal mechanism of STAMP 11.

Taken together, these findings provide a basic principle of peptide design and optimization to enhance the activity and specificity of narrow-spectrum antimicrobial agents. In our study, we successfully used a phage-displayed peptide to enhance the antimicrobial activity of STAMPs, while the mechanism of direct action on bacterial membranes positions peptide 11 as a potential candidate clinical treatment against *E. coli.*

## Conclusion

In our study, we attempted to use the traditional approach to optimize a STAMP 2 with activity against *E. coli* based on the principles of peptide chain length, amphipathicity, and charge number. A series of peptides were synthesized and tested for their anti-bacterial properties, hemolytic activity, cytotoxicity, and sensitivity. We found that STAMP is different from broad-spectrum antimicrobial peptides, and it is difficult to achieve a satisfactory effect by changing only one parameter. In contrast, we propose a feasible approach for the optimization of STAMP via the conjugation phage-displayed peptide, which enhances STAMP antimicrobial potency and stability. In our system, 11 showed enhanced narrow-antimicrobial activity against *E. coli* (TI_all_=0.028), while it had relatively high cell selectivity. Peptide 11 accumulated on the surface of the membrane by binding LPS and LTA of negative bacteria and positive bacteria, respectively, and permeabilized the outer membrane of Gram-negative bacteria, but it only induced depolarization of the cytoplasmic membrane in *E. coli*, disrupting the inner membrane integrity and eventually leading to target cell death. In summary, these results demonstrate a potential method for the design or optimization of STAMPs. Simultaneously, peptide 11 has potential as a novel agent against *E. coli* to help in preventing related diseases.

## Materials and Methods

### Bacterial strains

The bacterial strains *E. coli* ATCC25922, *E. coli* 078, *E. coli* K88, *E. coli* K99, *E. coli* 987P, *P. aeruginosa* ATCC27853, *S. agalactiae* ATCC13813, *S. typhimurium* C7731, *S. aureus* 25923, *S. aureus* 29213 and *S. epidermidis* ATCC12228 were obtained from the College of Veterinary Medicine, Northeast Agricultural University. *E. coli* UB1005 was kindly provided by the State Key Laboratory of Microbial Technology, Shandong University. *L. plantarum* 8014, *L. rhamnosus* 7469, *L. rhamnosus* 1.0911, *L. rhamnosus* 1.9205, *L. rhamnosus* 1.0385, and *L. rhamnosus* 1.0386 were obtained from the Key Laboratory of Food College, Northeast Agricultural University.

### Materials

Mueller-Hinton Broth (MHB) and Lactobacilli MRS Broth powder were obtained from AoBoX (China). Sodium dodecyl sulfate (SDS) was purchased from Sigma-Aldrich (China) and trifluoroethanol (TFE) was obtained from Amresco (U.S.A.). SDS and TFE were used after dilution to the desired concentration. Bovine serum albumin (BSA), *N*-phenyl-1-naphthylamine (NPN), 3,3-dipropylthiadicarbocyanine (diSC3-5), o-nitrophenyl-b-D-galactopyranoside (ONPG), Triton X-100, lipopolysaccharide (LPS) from *E. coli* 0111:B4, lipopolysaccharide (LPS) from *P. aeruginosa* 10, lipoteichoic acid (LTA) from *S. aureus*, 4-(2-hydroxyethyl)piperazine-1-ethanesulfonic acid (HEPES), ethanol (analytical grade, >99%), 3-(4,5-dimethylthiazol-2-yl)-2,5-diphenyltetrazolium bromide (MTT), tertiary butanol (analytical grade, 99%), acetone (analytical grade, 99%), glutaraldehyde (synthetic grade, 50% in H_2_O), propidium iodide (PI) and BODIPY-TR-cadaverine (BC) were purchased from Sigma-Aldrich (China). RPMI 1640, medium-high glucose (DMEM) and DMEM/F-12 and fetal bovine serum (FBS) were obtained from Gibco (China). Human embryonic kidney cells (HEK293T) and intestinal porcine enterocyte cells (IPEC-J2) were obtained from the College of Animal Science and Technology, Northeast Agricultural University (Harbin, China).

### Circular dichroism (CD) analysis

CD spectra were recorded at 25°C with a J-820 spectropolarimeter (Jasco, Tokyo, Japan). A 1.0 mm path length quartz cell containing a peptide (150 μM) solution was used along with 10 mM PBS, 30 mM SDS and 50% trifluoroethanol (TFE). At least three scans were acquired and averaged to improve the signal-to-noise ratio at 250-190 nm. The acquired CD spectra were then converted to the mean residue ellipticity using the following equation:

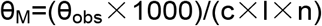

Where θ_M_ is the residue ellipticity (deg×cm^2^×dmol ^-1^), θ_obs_ is the measured ellipticity corrected for the buffer at a given wavelength (mdeg), c is the peptide concentration (mM), l is the optical path length (mm), and n is the number of amino acids.

### Determination of the minimum inhibitory concentration (MIC)

Pathogenic strains cells were cultured in Mueller-Hinton Broth (MHB), whereas beneficial strains were cultured in Lactobacilli MRS Broth (MRS). The MIC was determined in a 96-well plate. Briefly, logarithmic phase cultures of pathogenic strains and beneficial strains were diluted in MHB and MRS to a final concentration of 10^5^ CFU/mL, the series of peptides were serially diluted in 0.2% BSA solution, and the plate wells received aliquots of 50 μL each of the culture suspension followed by the addition of 50 μL of the diluted peptide; the final concentrations were 1, 2, 4, 8, 16, 32, 64, and 128 μM. Positive controls containing cells alone were incorporated. After incubation for 24-25 hours at 37°C, the optical density (OD) at 492 nm (Tecan, Austria) was measured, and the MIC was determined as the lowest concentration of peptide that resulted in inhibition of 95% of the bacterial growth. A minimum of three independent experiments (biological replicates) were conducted.

### Hemolytic activity test

The hemolytic activity of the peptides was determined according to a previously described method[49]. Fresh human red blood cells (hRBCs) were obtained from a healthy donor at the Hospital of Northeast Agricultural University. The hRBCs were pelleted by centrifugation and washed three times with PBS (1,000×g, 4°C, 5min). Then, the hRBCs were diluted 1:10 in 10 mM PBS (PH 7.4). Next, 50 μL of the hRBC solution and different concentrations of each peptide were mixed and incubated for 1 h at 37°C. The 96-well plate was centrifuged, and the supernatant was transferred to a new plate. Negative and positive controls for hemolytic activity were considered the hRBC suspension and hRBCs lysed with 0.1% Triton X-100 in PBS, respectively. Hemoglobin release upon lysis of the hRBCs was monitored at 570 nm using a microplate reader (TECAN GENios F129004; TECAN, Austria). The peptide concentration causing 5% hemolysis was considered to be the minimal hemolytic concentration (MHC). Hemolysis was calculated using the following equation:

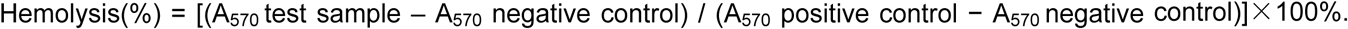

### Cytotoxicity assay

HEK293T and IPEC-J2 cells were used to assess the cytotoxicity of synthetic peptides by the MTT assay[50]. The cells (2.0×10^5^ cells/well in high-glucose DMEM or DMEM/F-12) were added to a 96-well plate, mixed in equal volumes with various concentrations of peptides (2-128 μM) and then incubated for 24 h at 37°C in 5% CO_2_. Cells without peptides served as controls. Next, the medium was replaced with 50μL of an MTT solution (0.5 mg/mL) and incubated for 4 h at 37°C. Subsequently, the formazan crystals were dissolved by the addition of 150 μL DMSO added to each well. The OD at 570 nm was observed using a microplate reader (TECAN GENios F129004; TECAN, Austria). The results were from three independent assays.

### Salt, Serum, Acid and Alkaline sensitivity

Each salt powder at a physiological concentration (300 mM NaCl, 9 mM KCl, 2 mM MgCl_2_, 16 μM ZnCl_2_, 12 μM NH_4_Cl, and 6 μM FeCl_3_) was dissolved in BSA stock solutions of polymer, and the subsequent steps were consistent with the MIC determination method. To evaluate the effect of serum, acid and alkaline conditions on antimicrobial activity, the peptides were incubated at three different serum levels (100%, 50%, 25%) and four different pH levels (pH=2, pH=4, pH=10, pH=12) for 4 h prior to MIC determination.

### LPS and LTA binding assay

LPS from *E. coli* O111:B4 and *P. aeruginosa* 10 and LTA from *S. aureus* were mixed with BC in 50 mL of Tris buffer (pH=7.4) and the solutions were incubated at room temperature for 4 h. Different concentrations of peptide in 50 μL were added to a 96-well plate after serial dilution. Then, a 50 μL aliquot of the LPS-probe mixture and LTA-probe mixture were added to each well. Subsequently, the fluorescence was measured (excitation λ=580 nm, emission λ=620 nm) on a spectrofluorophotometer (Infinite 200 pro, Tecan, China). Each test was performed independently in triplicate.

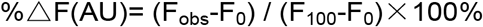

where F_obs_ is the observed fluorescence at a given peptide concentration, F_0_ is the initial fluorescence of BC with LPS (or LTA) in the absence of peptides, and F_100_ is the BC fluorescence with LPS (or LTA) cells upon addition of 20 μg/mL polymyxin B, which has a strong affinity for LPS and LTA as a positive control.

### Outer membrane permeability

*E. coli* 25922 and *P. aeruginosa* 27853 cells were suspended to 0.2 OD at 600 nm and incubated for 30 min in 5 mM HEPES buffer (pH 7.4, containing 5 mM glucose) containing 10 μM NPN. Subsequently, 100 μL of the cell suspension and 100 μL peptides of different concentrations (0.5-16 μM) were added to the 96-well plate. Fluorescence was recorded (excitation λ=350 nm, emission λ=420 nm with an F-4500 fluorescence spectrophotometer (Hitachi; Tokyo, Japan). Fluorescence was recorded until the fluorescence intensity remained constant. The values were converted to the percent NPN uptake using the following equation:

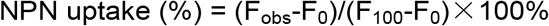

where F_obs_ is the observed fluorescence at a given peptide concentration, F_0_ is the initial fluorescence of NPN with *E. coli* and cells in the absence of peptides, and F_100_ is the fluorescence of NPN upon addition of 10 μg/mL polymyxin B.

### Inner membrane permeability

The ability of peptides to permeate the inner membrane of *E. coli* 25922 was assessed using cytoplasmic β-galactosidase with ONPG. *E. coli* 25922 cells, which were grown using MHB medium containing 2% lactose and centrifuged (5000 g, 5 min) to collect cells when the bacteria were in the mid-log phase. Then, the cells were suspended to 0.05 at 600 nm in 10 mM PBS (pH 7.4, containing 1.5 mM ONPG). Subsequently, 150 μL of bacterial culture and 50 μL of the peptide solution (final concentration of 2 μM) were added to a 96-well plate. The time-dependent effect of the peptides on ONPG fluorescence was measured at an absorbance wavelength of 420 nm.

### Cytoplasmic membrane depolarization

*E. coli* 25922, *P. aeruginosa* 27853 and *S. aureus* 29213 cells were grown to the mid-log phase at 37°C and diluted to an OD600 of 0.05 in 5 mM HEPES buffer (pH 7.4, containing 20 mM glucose). The cell suspension containing the 4 μM diSC3-5 was incubated for 1 h, and then, 100 mM KCl was added and incubated for 0.5 h. A 2 ml cell suspension was placed in a 24-well plate, and a peptide at a final concentration of 2 μM was added. The fluorescence was continuously measured for 800 s using an F-4500 fluorescence spectrophotometer (Hitachi, Japan) with an excitation wavelength of 622 nm and an emission wavelength of 670 nm, and the background fluorescence was measured.

### SEM characterization

*E. coli* 25922, *P. aeruginosa* 27853 and *S. aureus* 29213 cells were cultured in MHB at 37°C under constant shaking at 220 rpm until the logarithmic phase of growth and harvested by centrifugation. The precipitates were washed twice with 10 mM PBS and re-suspended to 0.2 OD at 600 nm. Then, bacterial cells were incubated at 37°C up to 60 min with peptides at a concentration of 2 μM. Subsequently, the samples were harvested (5000 g, 5 min) and fixed with 2.5% glutaraldehyde at 4°C overnight. The cells were dehydrated for 10 min in each of a graded ethanol series (50, 70, 90, and 100%). The cells have then transferred to a mixture (v: v = 1: 1) of 100% ethanol and tertiary butanol and absolute tertiary butanol for 15 min. Finally, the specimens were dehydrated in a critical point dryer with liquid CO_2_, and the dehydrated specimens were coated with gold-palladium and observed using a Hitachi S-4800 SEM (Hitachi, Japan).

### TEM characterization

Pretreatment with the bacterial samples was conducted in the same way as for SEM treatment. After treatment with a series of ethanol solutions (50, 70, 90, 100%) for 8 min, the samples have then transferred to a mixture (v: v = 1: 1) of 100% ethanol and acetone and absolute acetone for 15 min. Subsequently, the specimens were transferred to 1:1 mixture of absolute acetone and resin for 30 min and then to absolute epoxy resin overnight. Finally, the specimens were stained with uranyl acetate and lead citrate and observed using a Hitachi H-7650 TEM (Hitachi, Japan).

### Super-resolution microscopy (SRM)

*E. coli* 25922 cells were incubated in the presence of FITC-labeled peptide 11 at 2 μM at 37°C for 60 min. The mixture was centrifuged (1000 g, 5 min) and washed two times with PBS buffer. Then, the cells were resuspended and incubated with 10 μg/mL PI in PBS buffer for 15 min at 4 °C, and the extracellular PI dye was removed by centrifugation. A smear was created, and images were captured using a Deltavision OMX system with a 488 and 535 nm band pass filter for FITC and PI excitation, respectively.

### Flow cytometry

*E. coli* 25922, *P. aeruginosa* 27853 and *S. aureus* 29213 cells were diluted to 1×10^7^ cells ml^-1^ in PBS, and the peptide at a final concentration of 2 μM was incubated in the bacterial suspension at a PI concentration of 10 μg/ml for 60 min at room temperature. Images were obtained using a FACS flow cytometer (Bacton-Dickinson, USA) with a laser excitation wavelength of 488 nm.

### Statistical analysis

All data were subjected to one-way analysis of variance (ANOVA), and significant differences between means were evaluated by the Tukey test for multiple comparisons. The data were analyzed using the Statistical Package for the Social Sciences (SPSS) version 16.0 (Chicago, Illinois, USA). Quantitative data are presented as the means ± standard deviation (SD). P < 0.05 was considered statistically significant.

## Acknowledgments

This work was supported by the National Natural Science Foundation of China (31672434, 31472104, 31872368), and the China Agriculture Research System (CARS-35).

## Notes

The authors declare no competing financial interest.

## Author Contributions

P.T. and Z.L. contributed equally to this work, and they are both co-first-authors. P.T. and A.S. designed and conceived the experiments. P.T. and Z.L and Y.Z. conducted the main experiments assay. P.T. wrote the main manuscript text. C.S. and M.U.A and W.L. and X.Z. and A.S. supervised the work and revised the final version of the manuscript. All of the authors have read and approved the final version of the manuscript.

